## Supplemental figures for "*DMRT1* is a testis determining gene in rabbits and is also essential for female fertility"

^2^ École Nationale Vétérinaire d’Alfort, BREED; 94700, Maisons-Alfort, France.

^3^ Institute of Human Genetics, CNRS UMR9002 University of Montpellier; 34396 Montpellier, France.

^†^ and ^‡^ : These authors contributed equally to this work

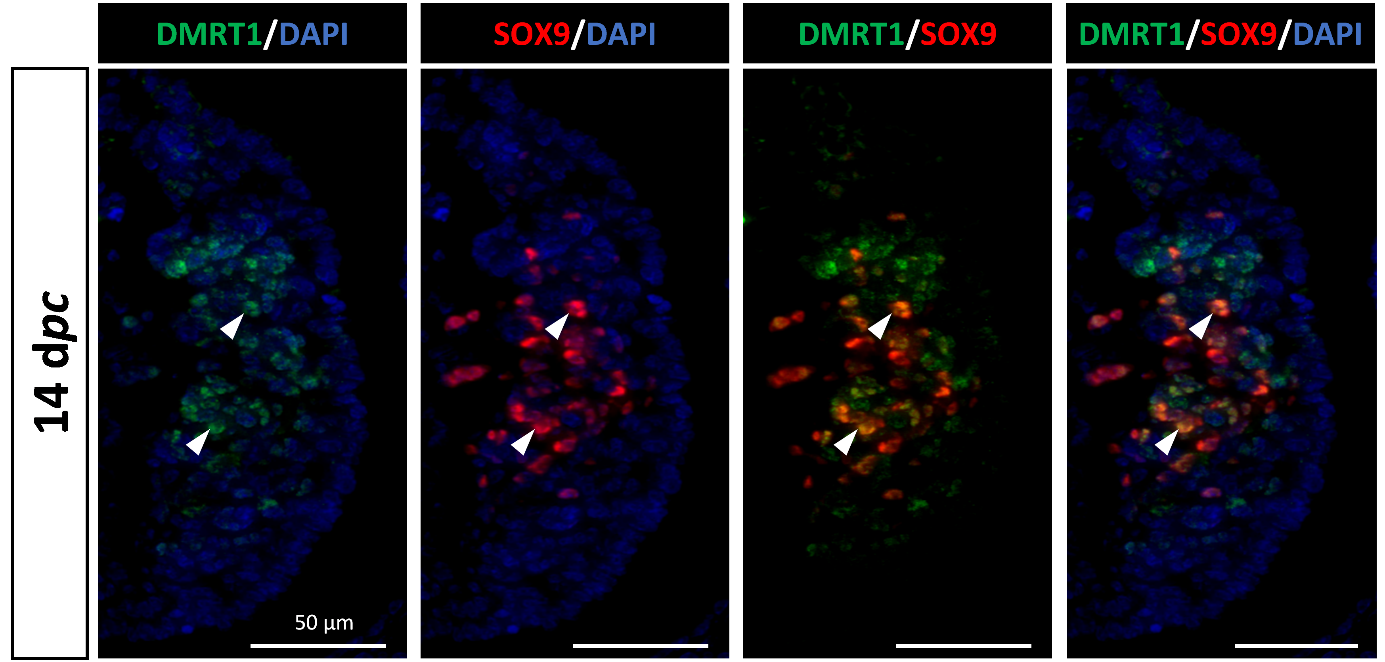
**Figure supplement 1. DMRT1 and SOX9 co-location on 14 d*pc* XY control gonad.** DMRT1 (green) and SOX9 (red) immunodetection in XY control gonad at 14 d*pc*. Nuclei were stained in blue (DAPI). Arrowheads: cells co-expressing DMRT1 and SOX9. Scale bar = 50 µm.

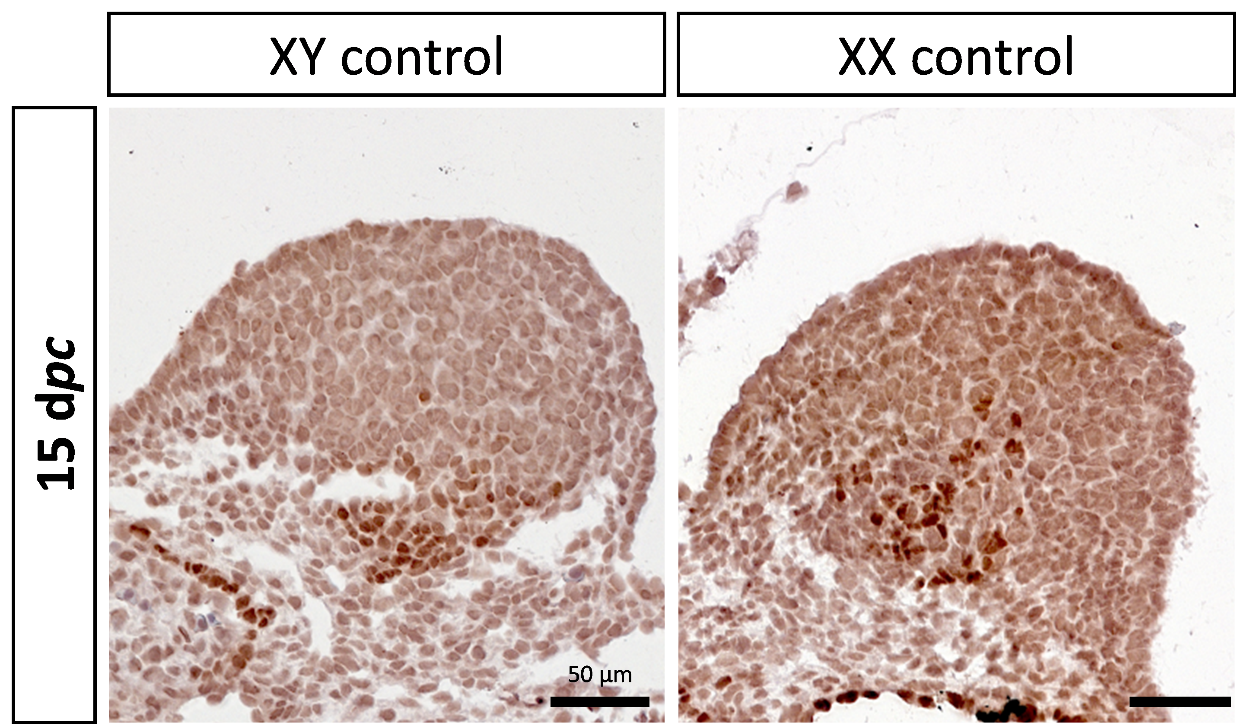
**Figure supplement 2. Identification of PAX8-positives cells in 15 d*pc* control gonads.** Immunostaining of PAX8 in XY and XX control gonads at 15 d*pc*. Scale bar = 50 µm.

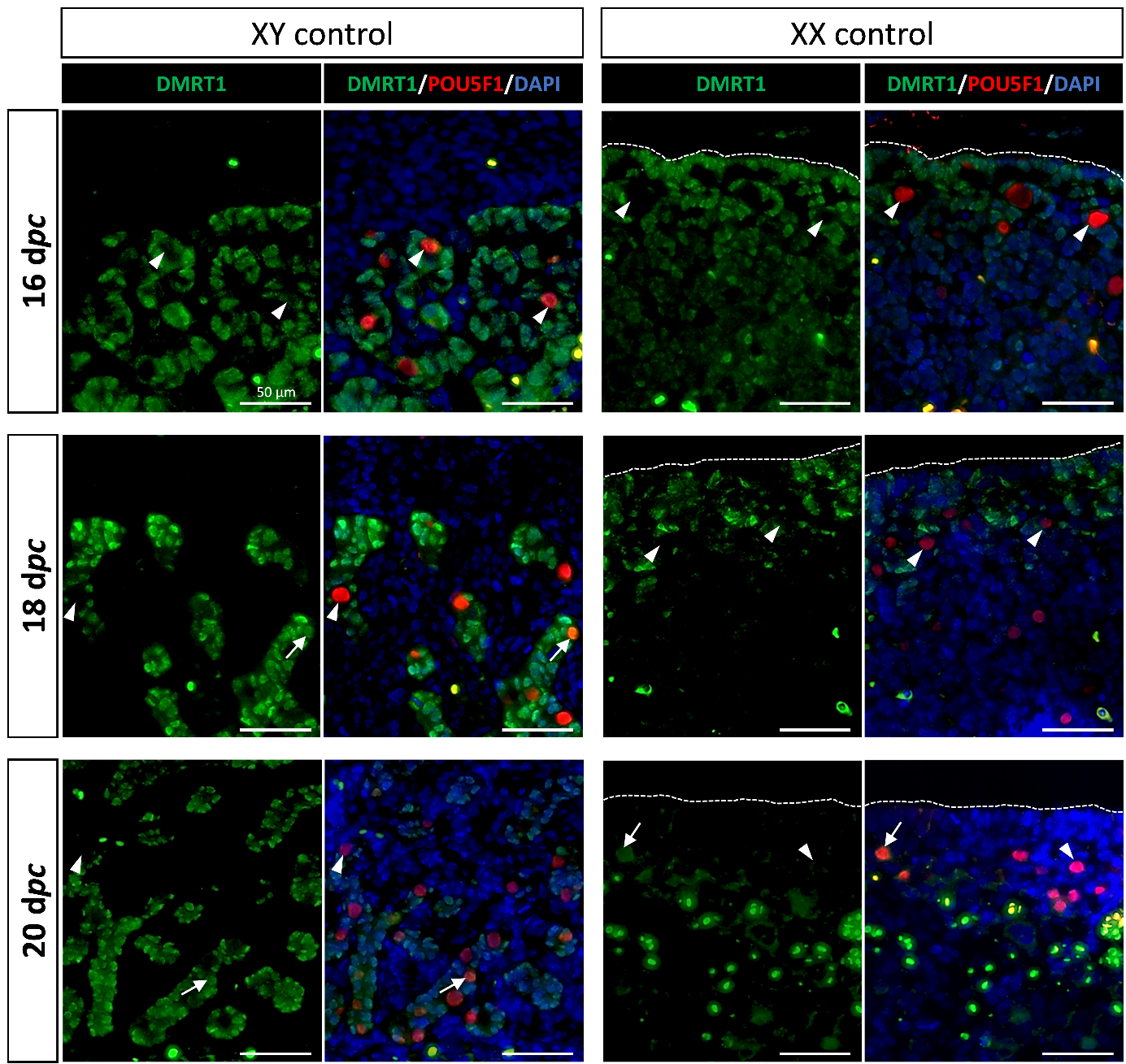
**Figure supplement 3. DMRT1 and POU5F1 co-detection in control gonads.** DMRT1 (green) and POU5F1 (red) immunodetection in XY and XX control gonads from 16 to 20 d*pc*. Nuclei were stained in blue (DAPI). Arrowheads: cells expressing POU5F1 only. Arrows: cells co-expressing POU5F1 and DMRT1. Dotted line: delimitation of the ovarian surface epithelium. Dots with intense green labeling: auto-fluorescence of red blood cells. Scale bar = 50 µm.

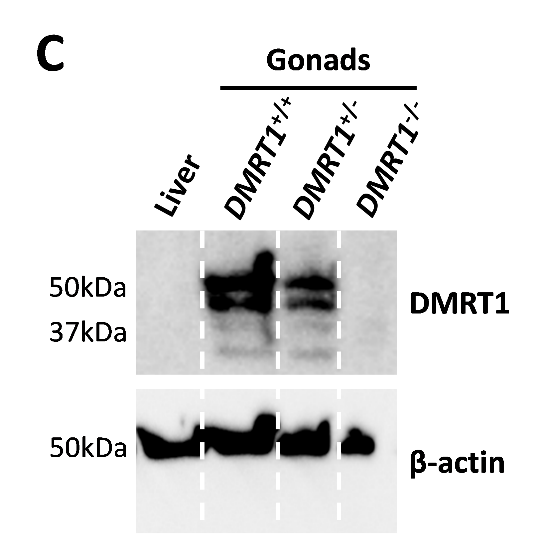

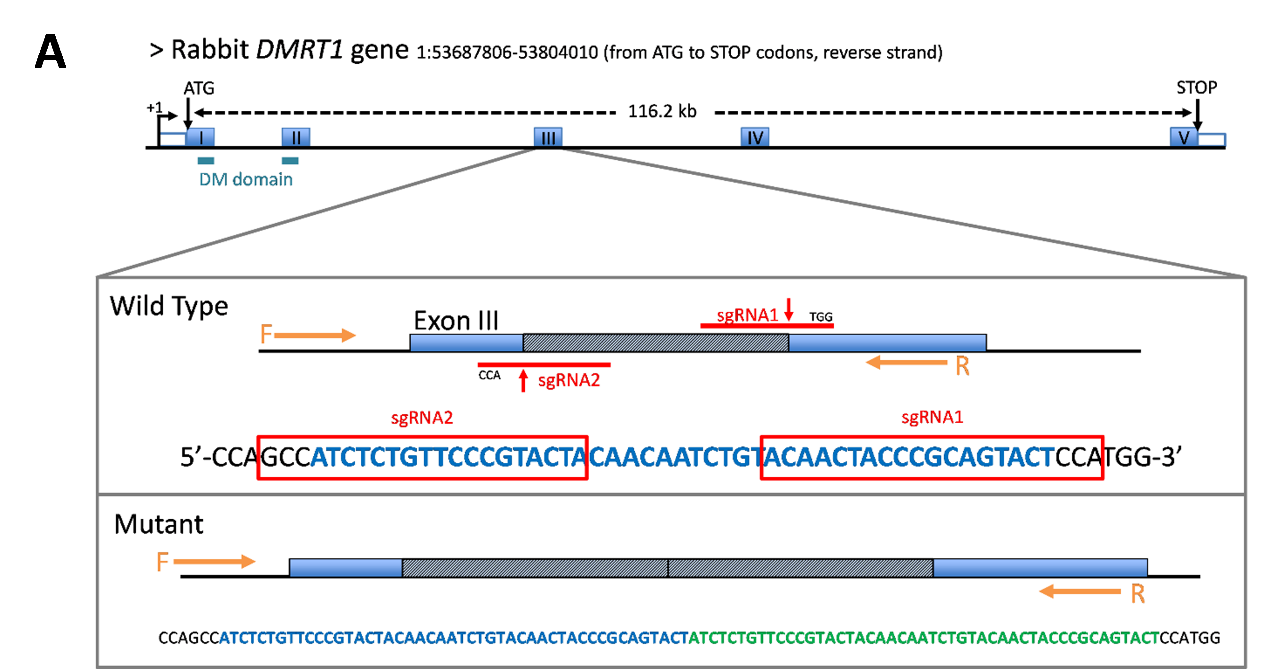

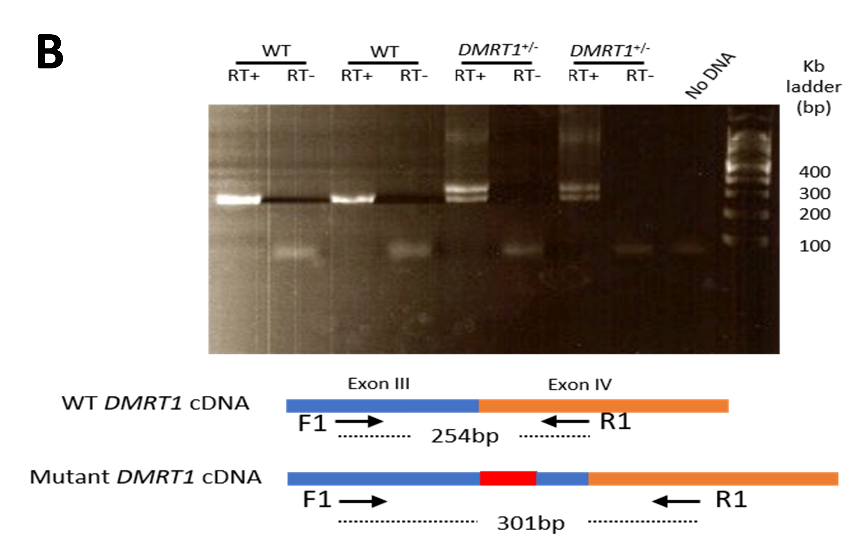
Figure supplement 4. *DMRT1* mutation using CRISPR/Cas9 in rabbits. (A) P osition of the two guides (sgRNA1 and sgRNA2) on the rabbit *DMRT1* transcript. Blue boxes represent the translated exons. The DM domain expands from exon I to exon II. The vertical red arrow indicates the Cas9-induced cleavage. The red box points to the sequence of the sgRNAs. The hatched box and bold blue letters point to the sequence between the theoretical and expected cleavage points. The cleaved fragment was inserted tandemly in the mutant allele at the cleavage site. Thus, by sequencing, a repeat was found (the repeats are written in blue and green bold letters). Consequently, the wild type and the mutant allele were characterized through PCR using the F/R set of primers and gel electrophoresis. +1: putative transcription start site; ATG: site of initiation of translation; STOP: stop codon. (B) Total RNA were reverse transcribed (RT+), and the amplified products were analyzed through gel electrophoresis. A unique amplicon of 254 bp was observed in PCR products from wild-type rabbits and two (254 and 301 bp) in PCR products from heterozygous rabbits. RT-: reverse transcription control (no reverse transcriptase). (C) Western blot with nuclear proteins extracted from the liver of control rabbits and from 7-8 gonads of 1-3 d*pp* *DMRT1^+/+^*, *DMRT1^+/-^* and *DMRT1^-/-^* rabbits.

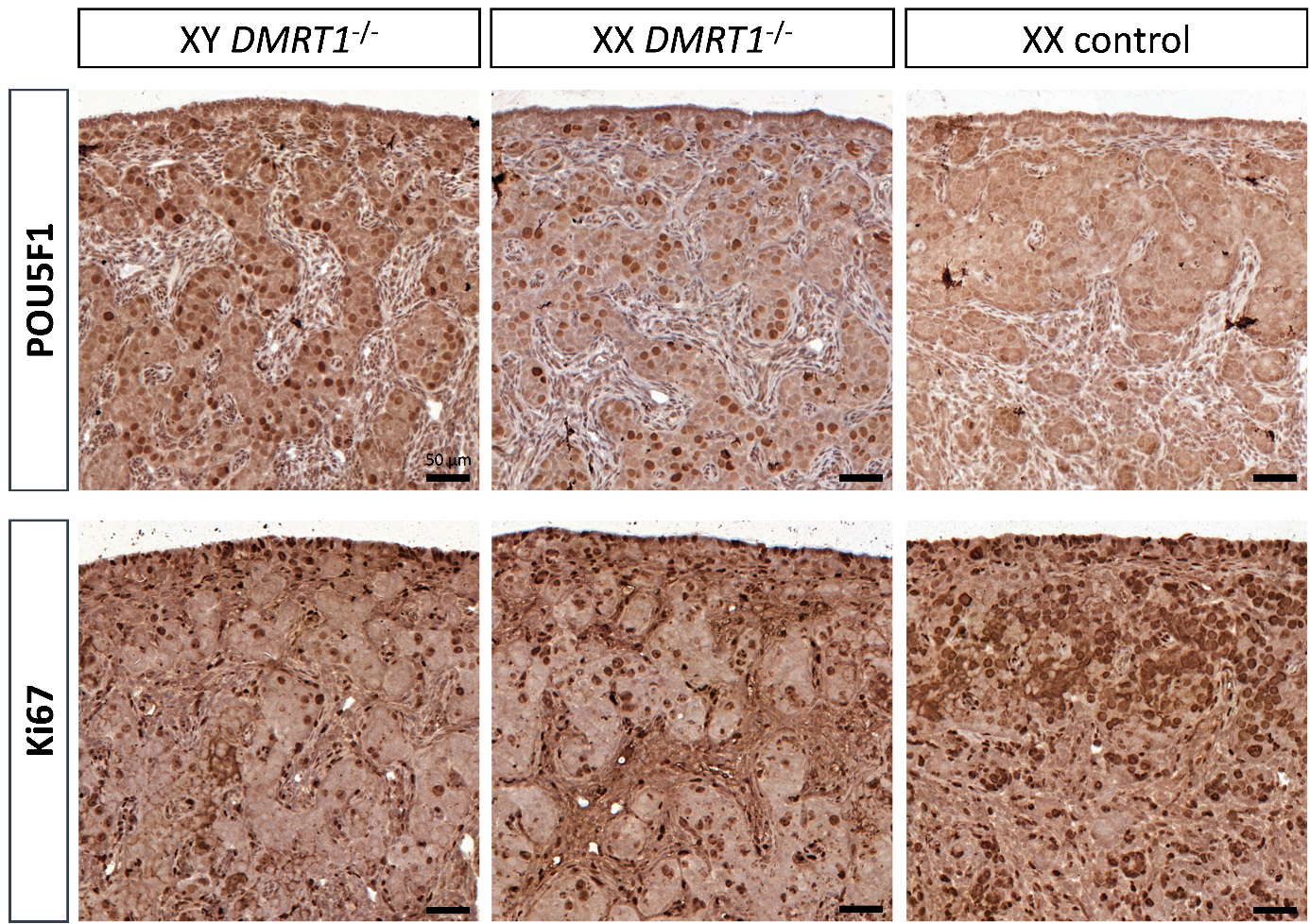
**Figure supplement 5.** **POU5F1 and Ki67 location on control and *DMRT1^-/-^* gonads at 3 d*pp*.** Immunostaining of POU5F1 (pluripotency marker) and Ki67 (a marker of the exit of the G0 phase of the cell cycle) on gonad sections from XY *DMRT1^-/-^*, XX *DMRT1^-/-^* and XX control at 3 d*pp*. Scale bar = 50 µm.

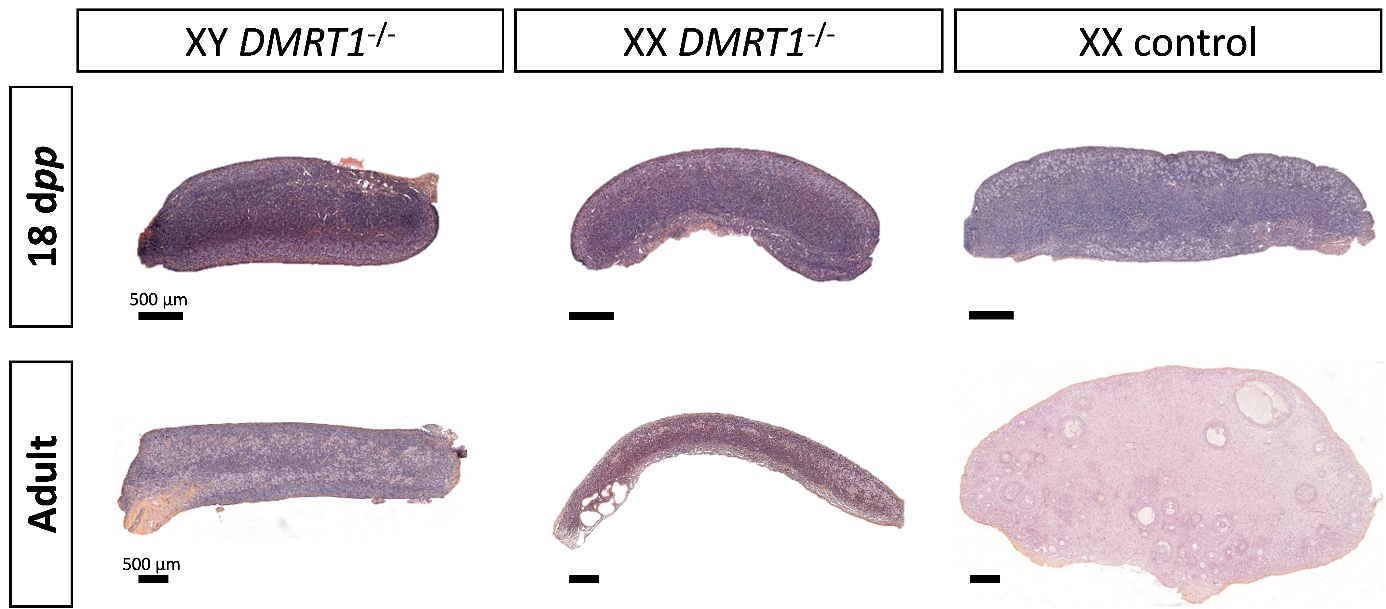
**Figure supplement 6.** **Evolution of gonadal size in XY and XX *DMRT1^-/-^* rabbits.** Hematoxylin and eosin staining of gonad sections from XY and XX *DMRT1^-/-^* gonads and XX control ovaries at 18 d*pp*, and in adulthood (4-9 months). Scale bar = 500 µm.

**Table supplement 1. List of DEGs in *DMRT1*^-/-^ gonads (XY and XX) compared to control gonads (XY and XX) (adjusted p-value <0.05 and |log2FC|>1).** List of deregulated genes (DEGs) between KO-XY *vs* Control-XY gonads (sheet 1), KO-XY *vs* KO-XY gonads (sheet 2), KO-XX *vs* Control-XX gonads (sheet 3), and Control-XY *vs* Control-XX gonads (sheet 4). The gene name of DEGs was based on their annotation or human homology (Craig et al., 2012).

**Table supplement 2. Clustering and expression values (TPM, Transcripts per million) of the 3460 DEGs.** Cluster membership of the 3460 DEGs with their expression data (TPM) according to the four genotypes (XY control, XY DMRT1^-/-^, XX DMRT1^-/-^, XX control).

**Table supplement 3. Primers used for genotyping PCR or RT-qPCR analyses.**

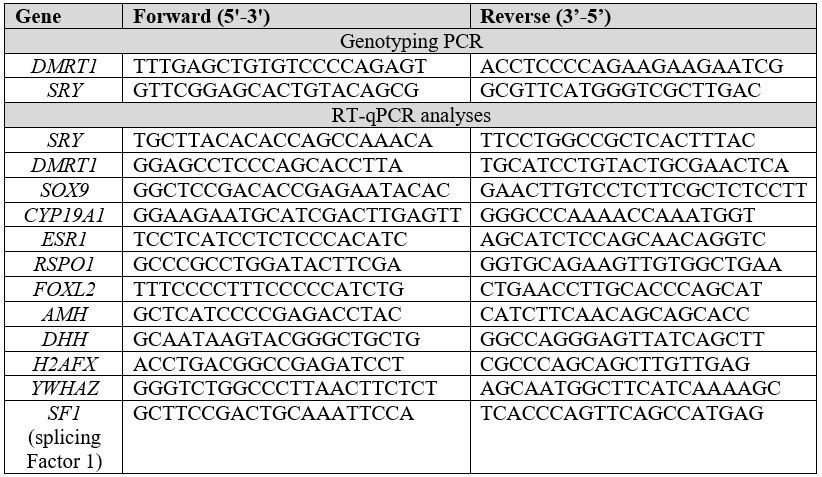

**Table supplement 4. Synthesized probes used for *in situ* hybridization.**

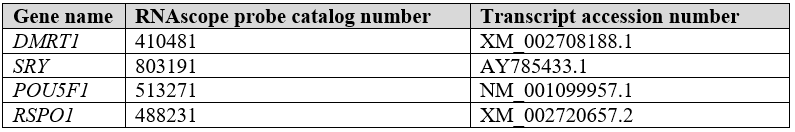

**Table supplement 5. List of antibodies used for immunohistochemistry (IHC), immunofluorescence (IF), or western blot (WB).**

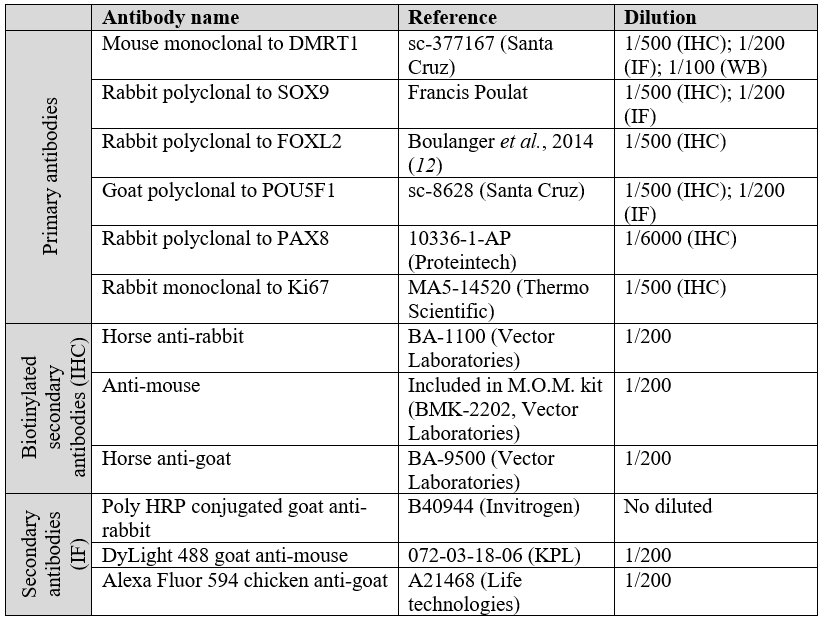
